# From hot to cold: dissecting lipidome adaptation in *Mycoplasma mycoides* and the Minimal Cell JCVI-Syn3B

**DOI:** 10.1101/2023.11.10.566608

**Authors:** Nataliya Safronova, Lisa Junghans, James P Saenz

## Abstract

Cell membranes insulate and mediate interactions between life and its environment, with lipids determining their properties and functions. However, the intricacies of how cells adjust their lipidome compositions to tune membrane properties remain relatively undefined. The complexity of most model organisms has made it challenging to characterize lipidomic adaptation. An ideal model system would be a relatively simple organism with a single membrane that can adapt to environmental changes, particularly temperature, which is known to affect membrane properties. To this end, we used quantitative shotgun lipidomics to analyze temperature adaptation in *Mycoplasma mycoides* and its minimal synthetic counterpart, JCVI-Syn3B. Comparing with lipidomes from eukaryotes and bacteria, we observed a universal logarithmic distribution of lipid abundances. Additionally, the extent of lipid remodeling needed for temperature adaptation appears relatively constrained, irrespective of lipidomic or organismal complexity. Through lipid features analysis, we demonstrate head group-specific acyl chain remodeling as characteristic of temperature-induced lipidome adaptation and its deficiency in Syn3B is associated with impaired homeoviscous adaptation. Temporal analysis uncovers a two-stage cold adaptation process: swift cholesterol and cardiolipin shifts followed by gradual acyl chain modifications. This work provides an in-depth analysis of lipidome adaptation in minimal cells, laying a foundation to probe the fundamental design principles of living membranes.

## Introduction

The cell membrane is a homeostatic organizational scaffold that insulates, while facilitating interactions of life with its surroundings. Ultimately, lipids determine the form and function of the cell membrane. Consequently, cells are continuously engaged in a balancing act to ensure that the composition of their membrane lipidomes is properly tuned to optimize both bioactivity and mechanical properties, such as bending rigidity^1^ and fluidity^2^. Although a tuneable membrane can be built from as few as two lipid structures^3^, cellular lipidomes are far more complex, ranging from 10s of lipids in bacteria^4^ to 100s in eukaryotic organisms^5678^. Despite the fundamental role of lipids in shaping membrane function and cellular fitness, how cells employ the collective properties of lipids to build responsive interfaces is still largely undefined.

To maintain optimal membrane property and function, cells must remodel their lipidomes in response to environmental changes that alter bilayer properties, such as temperature. Studying how cells manage their lipidomes in response to environmental change can provide critical insights for deciphering the principles underlying lipidome composition and regulation. For example, recent studies of lipidome adaptation in *Saccharomyces cerevisiae*^5^ and *Methylobacterium extorquens*^4^ revealed that phospholipid acyl chain unsaturation is regulated independently across different phospholipid head groups from bacteria to eukaryotes. The observation of head group-specific acyl chain remodeling prompted us to show that acyl chain structure has very different head group-specific effects on membrane fluidity, implicating a previously unrecognized lipidomic mechanism of physical homeostasis. Studies of the flexibility of lipid composition during adaptation have been dramatically enhanced by the emergence of quantitative shotgun mass spectrometry^9^, which has shone light on a previously inaccessible chemical space of cellular lipidomes. It is now possible to comprehensively characterize how lipidomes adapt to perturbations, Although there are still relatively few systematic studies of lipidome structure and responsiveness, the emerging body of work continues to reveal new patterns of lipidomic adaptation^108^.

The complexity of most cells poses a major challenge to understanding the principles underlying lipidome adaptation. Since many common model organisms such as *Saccharomyces cerevisiae* and *E. coli* have multiple membranes (e.g. plasma, ER, Nuclear, etc…, or inner and outer) with different lipidomic compositions, properties and functions, whole cell lipidomes provide limited insight into the behavior of specific membrane types. Purification of membranes is one solution to this problem; however, separating membranes is laborious and can result in considerable variations in purity. In this regard *Bacillus subtills* with its single plasma membrane is attractive. However, variations in the stereochemistry of acyl chain methylations that *B. subtilis* uses as a major mechanism of homeoviscous adaptation^1112^ cannot be resolved by shotgun mass spectrometry since they do not involve changes in lipid mass. Furthermore, the genomic complexity of most organisms provides a formidable challenge to characterizing and modeling the pathways involved in lipidomic sense and response^13^. Insights from more complex organisms are important for understanding the diverse biological functions that lipidomes support^14^. However, to probe the fundamental mechanisms of lipidomic architecture and adaptation, a minimal model organism, with a single plasma membrane and a lipidome that is amenable to shotgun mass spectrometry could provide valuable insight.

We propose *Mycoplasma mycoides subsp. capri str. GM12* and JCVI-Syn3B as minimal membrane model systems that bypass the challenges of commonly employed model organisms. Mycoplasmas are a class of relatively simple bacteria that hold promise as minimal living model membrane systems^15^. As notorious livestock pathogens, mycoplasmas have been studied for over a century and their potential as a minimal living membrane model was already recognized in in the 1960s^161718^. Recently, the genome of *M. mycoides* was reduced to the minimum set of genes that could support growth resulting in a Minimal Cell (JCVI-Syn3B)^1920^. Syn3B has the same core genome as *M. mycoides*, but lacks all non-essential genes^20^, providing a minimal counterpart for comparative studies. Understanding how simple organisms like *M. mycoides* and Syn3B compose and remodel their lipidomes can provide insight into fundamental lipidomic principles that are universal to diverse biological systems. However, the lipid composition of *M. mycoides* and Syn3B, and their lipidomic responses to environmental factors like temperature remain unexplored.

In this manuscript, we provide insights into the adaptation of *M. mycoides* and Syn3B lipidomes in response to varying growth temperatures. We took a systematic approach to explore lipidome composition and flexibility through an analysis of structural and global features^521^. Our findings reveal that certain lipidomic characteristics are conserved across organisms, from minimal cells to mammals. We discovered that head group-specific acyl chain remodeling is a key feature of temperature-induced lipidome adaptation. Notably, Syn3B’s reduced ability to remodel sphingomyelin is proposed to account for compromised homeoviscous adaptation. Additionally, cold adaptation is marked by a dual response: rapid changes in cholesterol and cardiolipin levels followed by more gradual acyl chain alterations. Our work represents the first detailed lipidome adaptation analysis in minimal cell lipidomes and opens a roadmap for exploring design principles of living membranes.

## Results

The goal of the present study was to understand how minimal bacterial cells adjust their lipidome composition when adapting to temperature change. To address this question, we used shotgun lipidomics to characterize the lipidome flexibility of two simple organisms, *M. mycoides* and Syn3B. We set out to examine the architecture of the lipidome in terms of the distribution of lipid abundances and structural features, and the lipidomic mechanisms of remodeling in response to temperature adaptation. We carried out growth experiments in 1 L continuous flow turbidostats, which allowed for continuous sampling at various temperatures, permitting us to characterize lipidome composition for adapted cells as well as at time intervals during adaptation. We thus characterized the lipid composition of cells in a temperature-adapted state, as well as during adaptation.

### 2.1 Establishing a viable temperature range

To determine a viable range of temperatures over which to observe adaptation, we first measured the growth characteristics of *M. mycoides* and Syn3B in batch culture. We characterized growth through measurements of cell density to construct growth curves from 25°C to the standard growth temperature of 37°C (Fig 1a). While *M. mycoides* grew across all temperatures, Syn3B growth was inhibited below 30°C. Growth rates estimated from an exponential fit of the growth curve data show that Syn3B growth rates are roughly 2-3 times lower than *M. mycoides* at a given temperature (Fig 1b). Although *M. mycoides* exhibits growth below 30°C, rates drop precipitously below this temperature. Maximum growth density also declines for *M. mycoides* below 30°C suggesting that both organisms grow optimally between 30 and 37°C (Fig. 1c). Based on these observations, we chose to examine the lipidomes of *M. mycoides* from 25 to 37°C, and Syn3B from 30 to 37°C.

**Figure 1.**
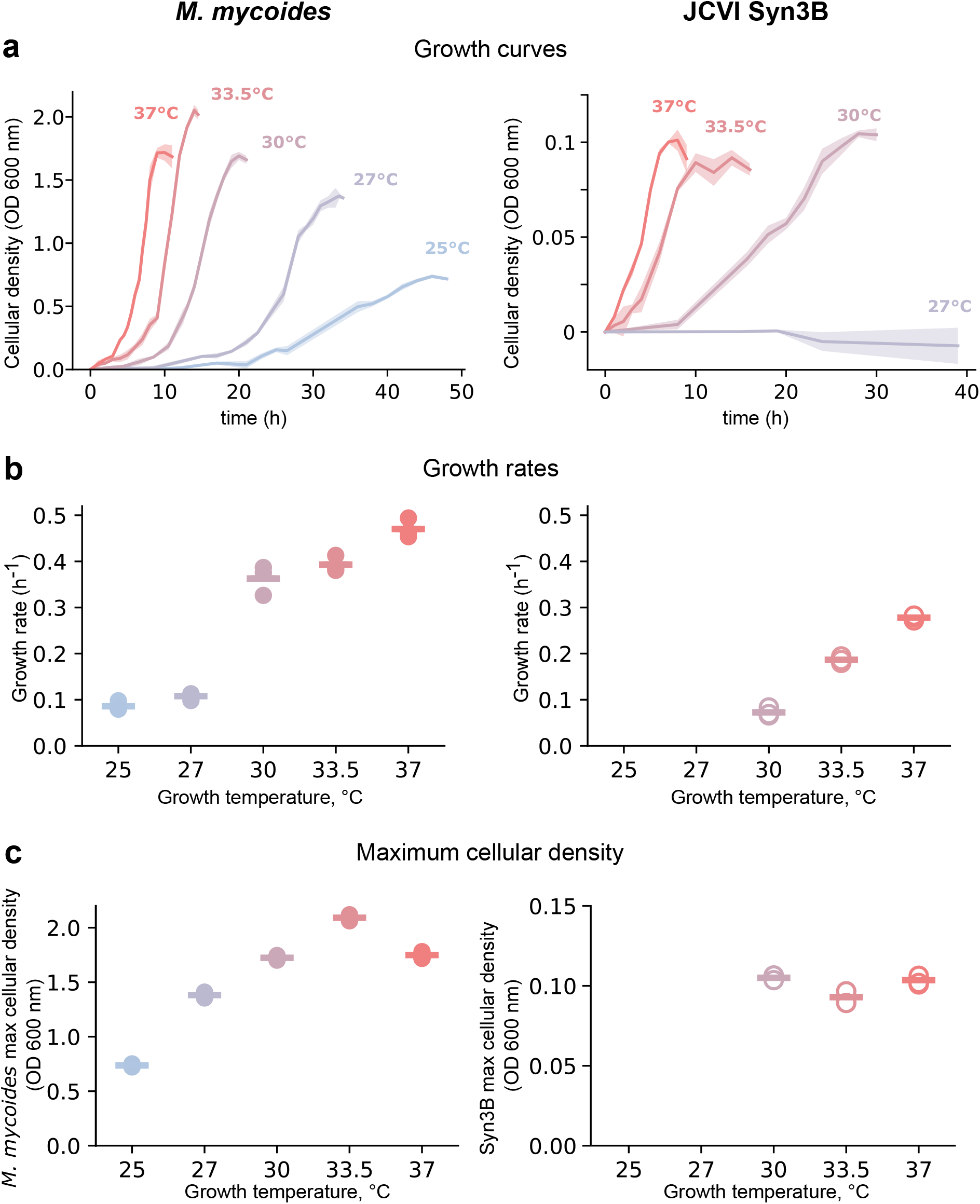
Mycoplasma growth analysis as a function of temperature. **a.** Growth curves of *M. mycoides* (left) and JCVI Syn3B (right) plotted against growth time (hours). Cellular density was recorded as bacterial sample absorbance at 600 nm. Each line represents mean +/- SD, standard deviations are shown as bands. n = 3. **b.** Bacterial growth rates, calculated from the growth curves shown in a for all growth temperatures. For growth rate calculations, see methods. mean +/- SD, n = 3.**c.** Maximum cellular density *M. mycoides* (left y-axis,filled circles) and JCVI Syn3B (right y-axis) reach at each of the growth temperatures examined. mean +/- SD, n = 3.

### 2.2 Lipidomes exhibit selective uptake from lipids provided in the media

Both *M. mycoides* and Syn3B have a restricted ability to synthesize lipids, and rely on uptake of lipids from the environment^182223^. In this study, we provided a complex lipid diet in the media by adding Fetal Bovine Serum (FBS), which contains a range of lipids including cholesterol, phosphatidylcholine and sphingolipids and free fatty acids (Supplementary Figure 2, (SP4 + FBS)). To determine whether the uptake of lipids from FBS is selective we compared the composition of FBS with lipidomes of cells from all conditions through principal component analysis (Fig 2a). Since both organisms are capable of synthesizing phosphatidylglycerol and cardiolipin from fatty acids taken up from the environment^1516242526^ (and Supplementary Figure 2), or scavenged from acyl chains of exogenous phospholipids^27^, we excluded these classes from the PCA analysis, only comparing across classes that are present in cells and FBS. The distribution of FBS lipids and cellular lipidomes demonstrates that *M. mycoides* and Syn3B have lipidome compositions that are distinctly different from FBS and from each other. This indicates that lipid uptake is selective. We next compared the lipidomes of *M. mycoides* and Syn3B across all temperatures to visualize how much they remodel their lipid composition during temperature adaptation (Fig.2b). This time we considered all lipid classes except for cholesterol-ester which is taken up, but enriched within the cell, and not in the membrane^2829^ (and Supplementary Figure 1b)). The distribution of samples shows that both organisms remodel their lipidomes at different temperatures, however, in different ways. Taken together, the results of our PCA analysis demonstrate that cellular lipidomes have distinct compositions from the FBS lipid source indicating selectivity, and that cellular lipidomes are remodeled at different growth temperatures.

**Figure 2.**
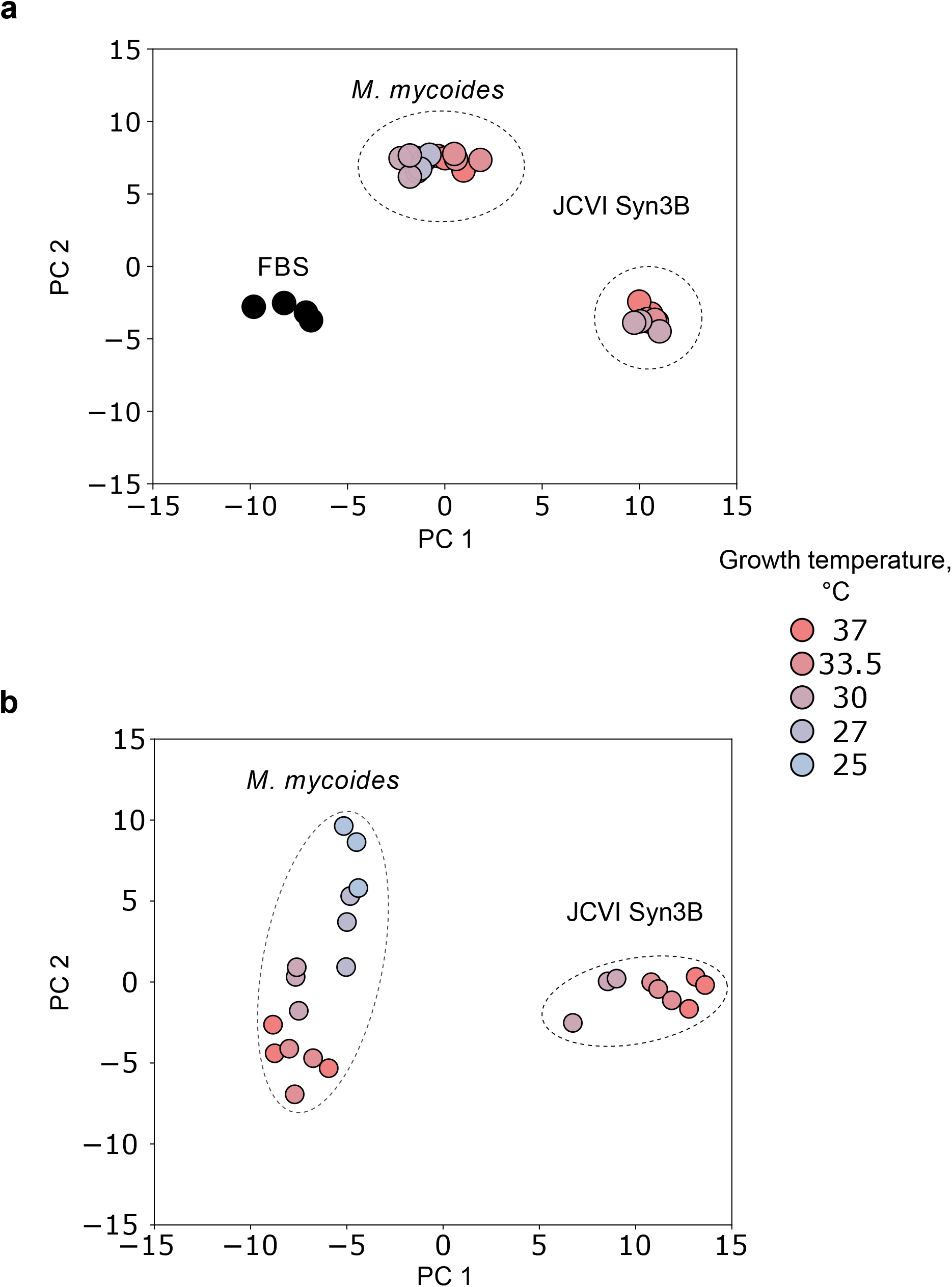
Principal component analysis shows the active mechanisms of mycoplasma membrane lipidome assembly. **a**. PCA score of membrane lipidomes of *M. mycoides* and Syn3B (bacterial lipidomes are labelled with dashed lines) at each of the growth temperatures examined and growth medium with FBS (black dots). Results for biological (for bacterial strains, n = 3) and experimental (for growth medium, n = 4) replicates shown. For the accuracy of direct comparison with FBS lipidome, PG and CL lipid classes were excluded from bacterial lipidomes prior the analysis. **b**. PCA score of membrane lipidomes of *M. mycoides* and Syn3B (distinguished with dashed lines), containing all lipid classes in mycoplasma lipidomes at all growth temperatures examined. n = 3.

### 2.3 Lipidome distribution and flexibility exhibit comparable features from minimal cells to mammals

Mycoplasmas are enveloped with remarkable membranes, which exhibit both eukaryotic and prokaryotic features; the eukaryotic sterol requirement for pathogenic strains is well-documented since the 1960s^30161731^ and is a rare feature among bacteria. At the same time, these bacteria synthesize their own cardiolipin^15^, which is typically a bacterial lipid, found mostly in mitochondria in eukaryotic cells^3233^. Are the fundamental principles of the mycoplasma membrane assembly similar to or different from other organisms? Understanding how variety of lipid inputs leads to organization of different types of cellular membranes is a crucial first step in unraveling the principles of lipidome organization. To this end, we explore several general lipidome features such as the distribution of lipid abundance, and the total change that the lipidome undergoes during adaptation to temperature. We compare these features against lipidomes from more complex bacterial and eukaryotic lipidomes, taking into consideration previously published lipidomes from a Gram-Negative bacterium (*Methylobacterium extorquens*)^4^, a eukaryotic microorganism (*S. cerevisiae*)^57^ and mammalian cell plasma membranes (isolated from rat basophilic leukemia cells)^34^.

We first explored how the abundances of individual lipid species within lipidomes are distributed, and how the distributions compare across organisms. Sorting the lipids in *M. mycoides* and Syn3B (Fig.3a, top) from the most to the least abundant immediately suggests the possibility of a power relationship between the lipidome size and lipid abundance. Moreover, lipidomes from a Gram-negative bacteria *M. extorquens* (whole cell lipidome), yeast (whole cell lipidome and isolated vacuole lipidome), as well as mammalian plasma membrane (isolated as GPMV from rat basophilic leukemia cells) (Fig.3a, bottom) appear to follow the same distribution.

Power law distributions are a characteristic of a vast variety of biological phenomena – ecosystems structure^35^, gene transcription regulation networks^36^ and, in particular, metabolic systems^37^. Hence, we postulate that this distribution might play a pivotal role in cellular membrane functionality. Furthermore, our study found an alignment of membrane lipid composition with the Pareto principle^38^, which empirically proposes that in many systems, 20% of inputs often account for approximately 80% of outputs. This principle seemed consistent across all samples examined (Fig.3b) except for the minimal cell. This suggests that, despite their unique chemical compositions, all living membranes might follow patterns seen in other natural processes. Moreover, this organizational principle remains consistent across different membranes, independent of lipid type, count, and organism complexity.

Examining the values of the power fit for different lipidomes revealed variations depending on the organism and specific conditions (Fig.3c). For *M. mycoides* and syn3B, the power fit becomes more negative at cooler temperatures, indicating that fewer species dominate a larger portion of the lipidome. This effect is subtler in *M. extorquens* and *S.cerevisiae*, especially when broader temperature ranges are considered. Interestingly, cholesterol-rich lipidomes of mycoplasma and the mammalian plasma membrane both exhibit a lipidome architecture characterized by a steeper, more negative slope in the power fit. In contrast, the internal membrane of the yeast vacuole has a less convex behavior. While the mechanistic implications of the power law distribution remain to be established, it appears that this is a conserved feature of lipidomic distributions that can vary as the membrane adapts to different growth conditions.

We next asked whether the amount of lipidomic remodeling required to adapt to temperature change is similar for different organisms. For example, do more complex organisms, or more complex lipidomes require less lipidome remodeling to adapt to a given change in temperature? The extent of lipidome remodeling that occurs between two conditions (e.g. high and low growth temperature) can be quantified by summing up the absolute change of each lipid, to yield the total lipidomic change (Supplementary Figure 2(b)). As expected, the change in total lipidomic remodeling increases as the change in growth temperatures increases. When we normalize total lipidomic remodeling to estimate the mol% change per degree Celsius the variation across the four organisms considered spans a range from roughly 2 to 3 mol%/C (Fig.3d). Therefore, the amount of lipidome remodeling required for temperature adaptation from minimal cells to mammals falls within a ∼70% range of variability. Thus, it appears that the amount of lipidomic remodeling required for temperature adaptation, at least within mesophilic temperature ranges, is constrained within a less than two-fold range of variability.

### 2.4 Lipidome composition of M. mycoides and Syn3B

Lipidomes are made up of complex mixtures of lipids, each possessing a unique combination of structural features that determine how they contribute to the physical and chemical properties of the membrane. For example, phospholipid acyl chains can be different lengths, or have varying numbers of double bonds, both of which will influence the properties of the membrane in different ways; longer acyl chains result in thicker^39^ and less fluid membranes^10^, whereas more double bonds result in more fluid membranes^404142^. One way to make sense of the structural complexity of the lipidome, is by considering the distribution of biophysically relevant structural features, rather than individual lipid species. In this sense the lipidome can be aggregated by lipid structural features, such as class (e.g. sterols and phospholipids) or acyl chain structure (e.g. chain length and number of double bonds). Through this lens, it is possible to reveal patterns of lipidome adaptation that are otherwise hidden in the complex milieu of individual lipids. This approach has previously revealed how different phospholipid head groups have distinct acyl chain configurations that are independently regulated during adaptation^45^. Here we dissect the lipidome in terms of its aggregated lipid features to reveal the underlying patterns of lipidome composition in *M. mycoides* and Syn3B.

At the lipid class level, the lipidomes of *M. mycoides* and Syn3B contain a sterol (e.g. cholesterol), glycerophospholipids (e.g. phosphatidylglycerol – PG, Cardiolipin – CL, phosphatidylcholine – PC) and phosphosphingolipids (sphingomyelin – SM) (Fig 4a). Both lipidomes are dominated by cholesterol, which accounts for at least half of the total lipid abundance. Similarly high sterol content in mycoplasmas has been previously reported^43^. In the absence of a cell wall, cholesterol strengthens the membrane barrier function and modulates mycoplasma membrane physical properties^4445^, regulates membrane fluidity^15^ and possibly aids in mycoplasma-eukaryotic plasma membrane interactions^46^. The presence of PG and cardiolipin, which are internally synthesized by both *M. mycoides* and JCVI Syn3B (Supplementary Fig.2), corroborates the previous studies on mycoplasma phospholipids^1615242526^. The synthesis of PG and CL requires an exogenous source of fatty acids. The average abundance of PG and CL is the comparable in both strains, suggesting synthesis and regulation of these lipids is conserved in the minimal cell. PC and SM which cannot be synthesized by *M. mycoides* or Syn3B are taken up from FBS in the media^29^. SM levels at around 10 mol% (*M. mycoides*) and 20 mol% (syn3B) are in the range identified for various eukaryotic membranes, depending on the tissues and characterization method^47218^. The considerable difference in the relative abundance of PC and SM taken together with higher cholesterol levels in the minimal cell represent the largest difference between *M. mycoides* and JCVI Syn3Blipidome architecture at the class level.

**Figure 3.**
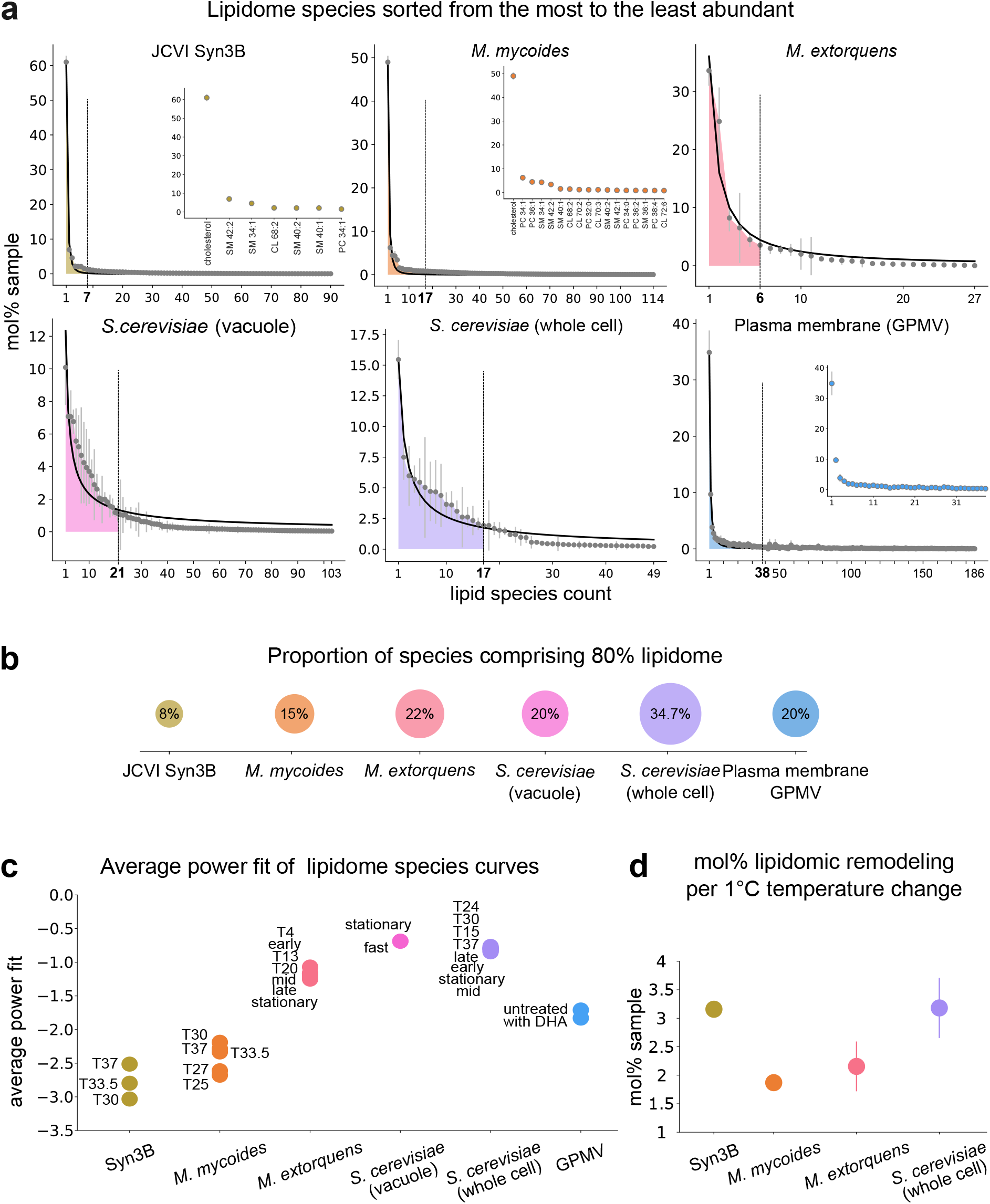
Cellular membrane lipidomes share fundamental organization principles. **a.** Lipidome species of: Syn3B and *M. mycoides*; *M. extorquens*^4^; *S. cerevisiae* vacuole^7^; *S. cerevisiae* whole cell lipidome^5^; plasma membrane GPMV, from RBL cells^34^. Scatter plots represent averages across several growth conditions available, with variability shown as error bars. For mycoplasma lipidomes, all growth t°C were considered. For other organism lipidomes, conditions included are listed in methods. Black lines represent the average power fit to the species distribution. Filled areas under lipidomes with a cutoff lipid number show 80% of the total lipid distribution. Insets for Syn3B, *M. mycoides* and GPMV show the species comprising 80% lipidome abundance on the main plot. **b**. Percentage of lipid species in lipidomes that account for 80% of the total abundance in each of the organisms, shown in **a. c**. Power fits for all growth conditions, considered in **a**, shown separately for each organism,with respective labels. **d**. Mol% total lipidomic remodeling per 1°C change in growth t°C, calculated for organisms, where temperature conditions are available.

**Figure 4.**
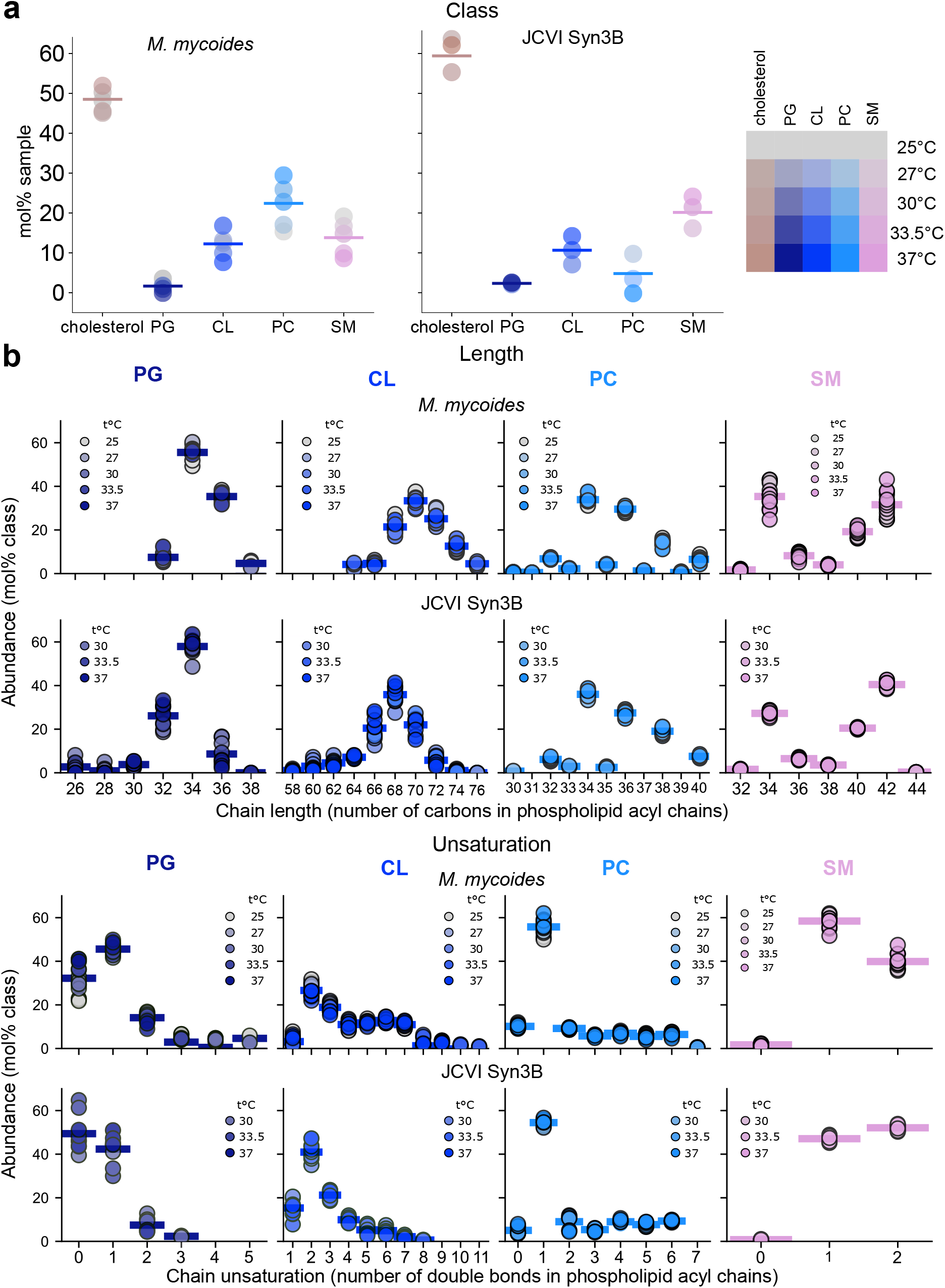
Lipidome composition of *M. mycoides* and Syn3B. **a.** Lipid class distribution in mycoplasma lipidomes grown at different temperatures: *M. mycoides* (left) and JCVI Syn3B (right). Each dot shows mol% abundance average for 3 biological replicates for all growth temperatures. Total average across all growth temperatures is shown as a bar line. **b**. Acyl chain features distribution in each phospholipid class, introduced in **a**. Features are split to show both length (left) and unsaturation (right) profiles for *M. mycoides* (top) and Syn3B (bottom) at different growth temperatures. Sample dots represent temperature averages (n = 3), with bar lines showing the total average across all growth temperatures.

The lipidome can also be aggregated according to the acyl chain features of each phospholipid class (e.g. PG, CL, PC, and SM) (Fig 4b). Acyl chains can be characterized by the number of double bonds (unsaturation) or chain length. Acyl chain length and unsaturation profiles are comparable for *M. mycoides* and Syn3B with the exception internally synthesized PG and CL chain length, which is slightly shorter in Syn3B. Total length of 34 carbons is the most abundant in phospholipids with 2 acyl chains (PG and PC), followed by length of 36 carbons. This is in accord with glycerophospholipids acyl chain profile found in yeast^5^ and rat basophilic leukemia cells^34^ (RBL); in nonpolarized MDCK cells GPL length distribution is reversed, but nevertheless 34 and 36 carbons remain dominant^21^. Total unsaturation of 1 double bond per lipid is prevalent in mycoplasma (with the exception of Syn3B PG), which is also the case in yeast^5^, MDCK^21^ and RBL cells^34^. Moreover, the sphingomyelin acyl chain profile of mycoplasma is very close to that of other published mammalian lipidomes, with length ranging between 32-44 carbons and unsaturation predominantly distributed between 0-2 doble bonds per lipid^2134^. These observations illustrate how mycoplasma utilizes mammalian lipids to construct a membrane with acyl chain features very close to those of mammalian cell, while constructing a structurally unique lipidome. How do mycoplasmas employ their distinct lipid membranes to adapt to environmental perturbations?

### 2.5 M. mycoides and Syn3B exhibit homeoviscous adaptation

One of the most well-established principles in membrane biology is homeoviscous adaptation, whereby cells modulate their lipidome composition in response to changing temperature in order to maintain optimal membrane fluidity^48^. Homeoviscous adaptation is conserved in organisms from microbes to mammals^41^, however it is not known whether minimal organisms like *M. mycoides* and Syn3B exhibit or require homeoviscous adaptation. Our observations showing lipidomic remodeling across varying temperature suggests that despite their simplicity, *M. mycoides* and Syn3B are capable of homeostatic regulation of their membrane’s physical properties. To address this, we probed membrane fluidity across varying growth temperatures, to establish whether homeoviscous adaptation is conserved in minimal cells.

To monitor membrane fluidity we employed a recently developed fluorescent membrane fluidity probe called Pro12A which is derived from the solvatochromatic dye c-laurdan ^4950^. The emission spectra of Pro12A is sensitive to the hydration of the bilayer, which in turn is tightly coupled with lipid order and membrane fluidity. By taking the ratio of two emission maxima, a general polarization (GP) index can be calculated, in which higher values correspond to lower fluidity. An advantage of Pro12A over c-laurdan is that because of its bulky polar head group, it selectively labels the outer leaflet of the plasma membrane, thereby preventing labeling of non-membranous hydrophobic compartments within the cell^49^. Pro12A therefore makes it possible to selectively monitor membrane fluidity in situ through bulk spectroscopic measurements, which is essential for small mycoplasma cells that challenge the resolution limits of microscopic analysis. The lipid order of *M. mycoides* and Syn3B membranes reported as Pro12A GP values ^4950^ are summarized on Fig.5a. On average, Pro12A GP of live mycoplasma cells are in the range of ∼0.35-0.45, which is a relatively high lipid order (low fluidity) and comparable with GP values for mammalian plasma membranes^49^. Syn3B membranes are more ordered across the temperature range, than *M. mycoides*, which is consistent with higher cholesterol and SM content in Syn3B (Fig.4a).

The capacity for homeoviscous adaptation can be assayed by measuring the change in lipid order across different growth temperatures. We would expect lipid order (GP values) to remain constant if homeoviscous adaption were active. In the absence of homeoviscous adaptation, lipid order would increase with decreasing temperatures, and approach the change observed for non-adaptive liposomes, which in this case we have reconstituted from pure phospholipid and cholesterol. We plot the change in lipid order as the delta-GP for a given temperature range in Figure 5b. The change in lipid order for 37°C- 30°C temperature difference for *M. mycoides* and the minimal cells is about 3-fold lower, than that for the synthetic liposomes GP (Fig.5b), which is an indication that both bacterial strains effectively maintain their membrane lipid order in a narrow range, when compared to non-adaptive membranes ^48^. The two bacterial strains show differences in their adaptation to temperature: *M. mycoides* appears to be more efficient in restraining its membrane order to a narrower range across large growth temperature span, than Syn3B (Fig.5b). These observations demonstrate the presence of homeostatic sense and response mechanisms that allow *M. mycoides* and Syn3B to control their membrane biophysical state when challenged with changing growth temperature. Moreover, the data suggest that the *M. mycoides* might possess a more robust mechanism for adaptation than Syn3B. Nonetheless, homeoviscous adaptation is conserved even in a minimal cell membrane.

**Figure 5.**
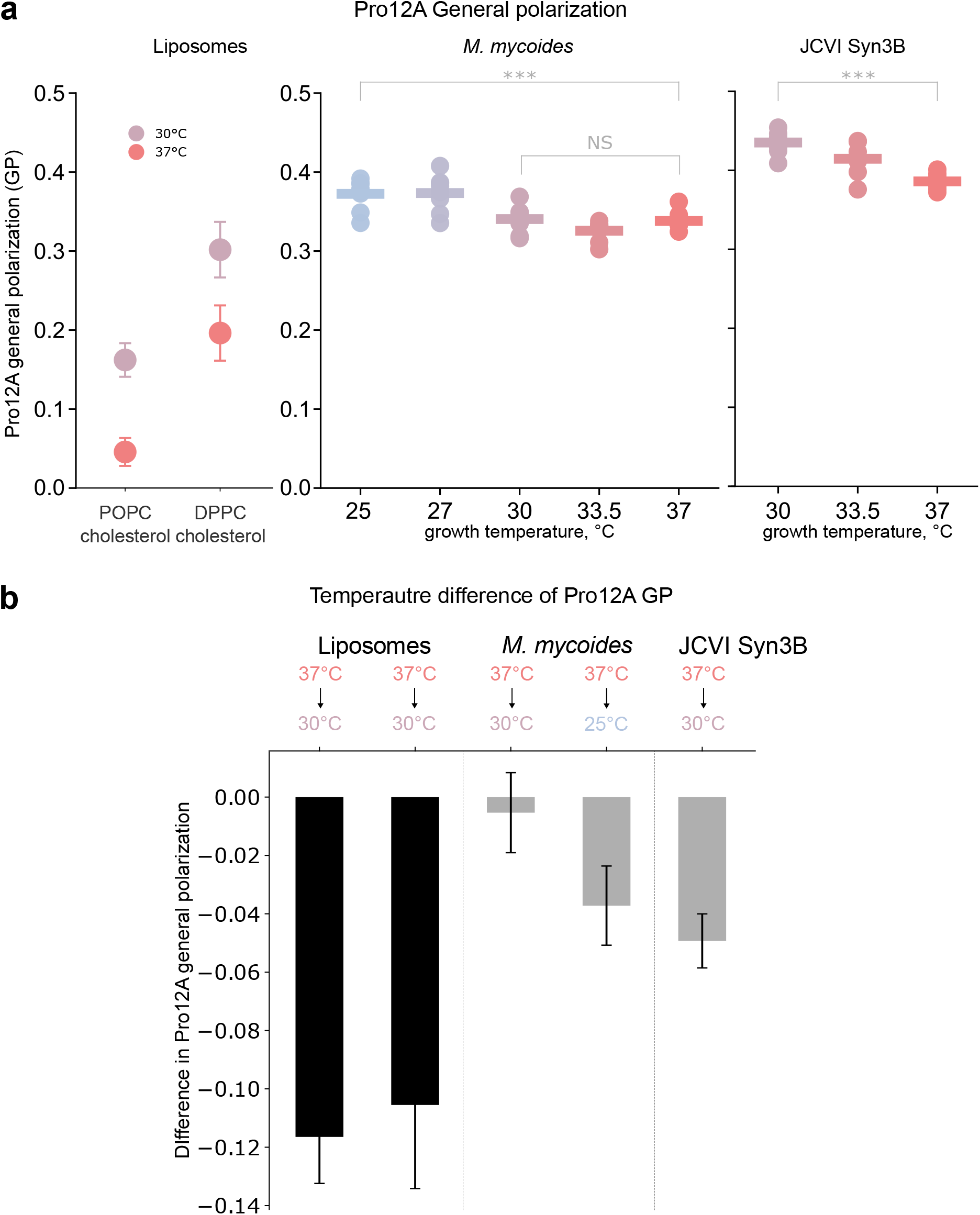
Mycoplasma membranes homeoviscously adapt. **a.** Pro12A general polarization (GP) values for *M. mycoides* and JCVI Syn3B membranes at each of the growth temperatures examined. Data for 3 biological and 9 analytical replicates are shown for each sample, with colored lines indicating the average. Gray lines show overall GP average for each bacterial strain across all conditions measured. The corresponding GP values of synthetic liposomes (left) are shown as a reference point for non-adaptive liquid-ordered membranes temperature response; mean +/- SD, n = 9. Mann-Whitney Ranking U-test results are shown for bacterial GP values at 30°C- 37°C (both *M. mycoides* and Syn3B) and 25°C-37°C (*M. mycoides*) **b.** The respective GP differences in 37°C- 30°C and 37°C-25°C shown for bacterial (gray bars) and synthetic (black bars) membranes to showcase the evidence for homeoviscous adaptation in mycoplasma.

### 2.6 Mechanisms of lipidomic remodeling

Having demonstrated that both *M. mycoides* and Syn3B can remodel their lipidomes to homeostatically maintain membrane fluidity, we next investigate which elements of the lipidome contribute to adaptation. We start by inquiring how the lipidome varies at the species level, and then evaluate variation in terms of the lipid features that we introduced in **figure 4**.

#### 2.6.1 Lipid species variability

In general, variations in individual lipids across growth conditions tend to be on the order of fractions of fold-change, unlike, for example, variations in cellular mRNAs which can exhibit several-fold change^5152^. Indeed, the average mol% change in lipid abundance for *M. mycoides* and Syn3B between 30 and 37°C is about 0.14 mol% and 0.25 mol%, respectively. But how many lipids are highly variable and what fraction of lipidome variability do they account for? To examine how high and low abundance lipids contribute to total variability, we took a simple approach to split the lipidome into two groups of high and low abundance. The distribution of lipid species abundances has a single point of maximum curvature that occurs at around 0.5-1 mol% average lipid abundance (Fig.6a). This point visually separates the lipidome into two distinct ‘tails’ of high abundance (on the left) and low abundance lipids (on the right). The presence of the maximum curvature point indicates a transition in the abundance distribution behavior from the few dominant high-abundance lipids to the bulk of low-abundance species. To precisely locate this point on lipid distribution curves, we searched for the maximum curvature using Kneedle algorithm, originally developed by Satopää et.al^53^. The dashed line on Fig.6a shows the resulting curvature point. The line separates 4-5 most abundant species from the rest of the lipidome. The inset bar on Fig.6a shows that these species sum up to about 68-75 mol% sample, which is very close to Pareto distribution (Fig.3a,b). Is lipid species variability determined by which abundance ‘tail’ they belong to?

When we plotted lipid species variability against mean abundance (Fig.6b), we observe that most lipids vary less than 0.5 mol% and that only those assigned as high-abundant (shown in color on Fig.6b) exceed 1 mol% change (Fig 6b). In general variability is scaled non-linearly with abundance (decreasing with abundance), similar to the distribution of abundance (Fig 3). The most abundant species are remodeled in a distinct way from the rest of the lipidome, while the rest of the lipidome followed a similar remodeling trend below 1 mol% change for all species. The most abundant and variable (in absolute terms) species are relatively conserved in mycoplasma – besides cholesterol, it is mainly SM (in particular, 34:1 and 42:2 species), followed by the 1-2 species from the second most abundant phospholipid in the lipidome (PC – for *M. mycoides*, CL – for Syn3B). Strikingly, the most variable lipids only account for around half of the total lipidomic remodeling (Fig. 6c) or even less (for Syn3B). Therefore, it appears that half the lipidome adaptation is due to small variations in many low abundance lipids.

**Figure 6.**
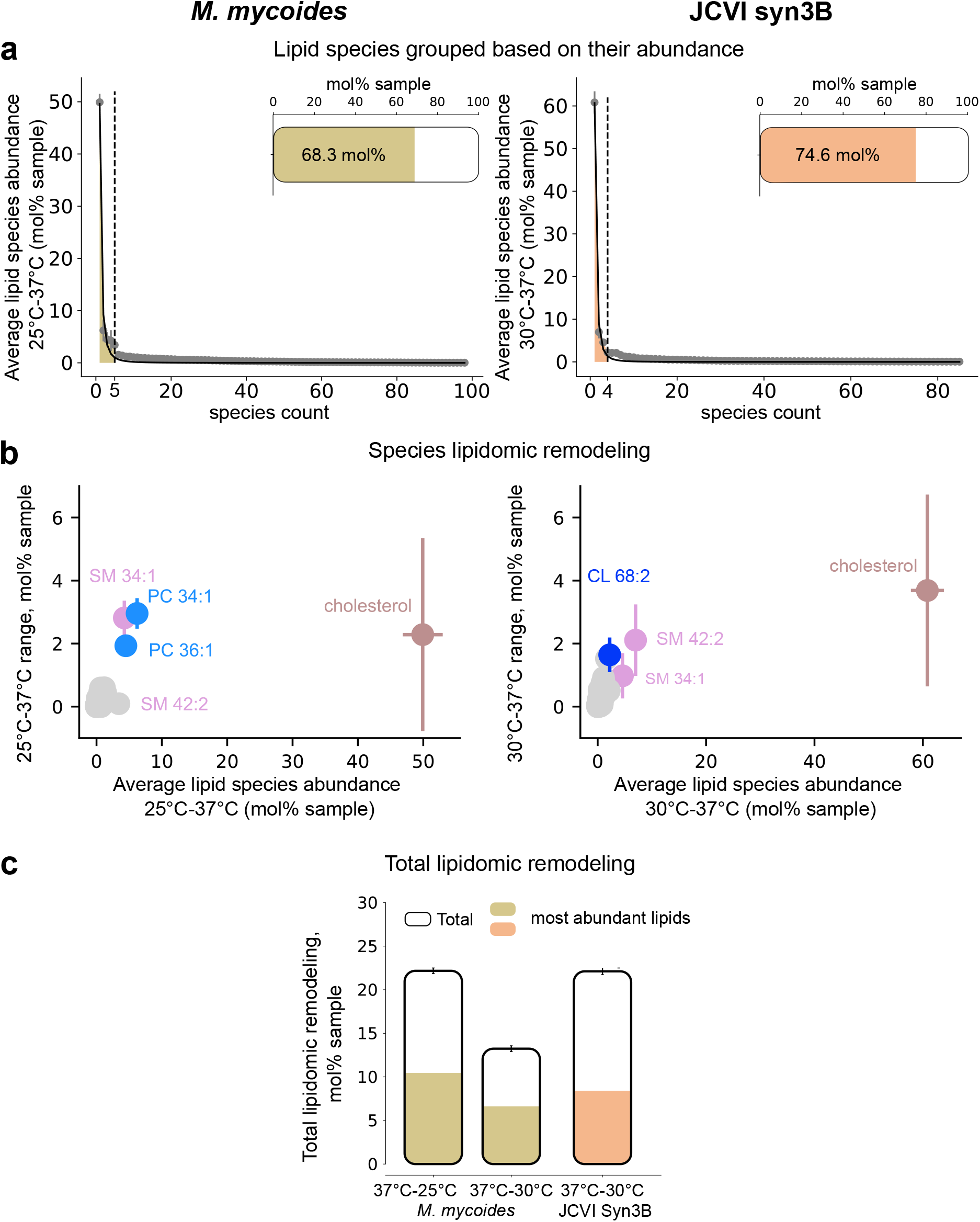
Mechanisms of lipidomic remodeling in Mycoplasma. **a.** Species average abundance at 37°C and 25°C (*M. mycoides*) and 37°C and 30°C (Syn3B) growth temperatures sorted from largest to lowest abundance and split into two functional abundance groups: largest abundance and low abundance lipids. The dashed line shows the split point determined by the steepest curvature of the abundance curve. The largest abundance lipid fraction is further highlighted in color. The bar at the top right of each abundance curve shows the mol% abundance taken by the large abundance fraction in total sample. **b**. Species absolute abundance change with temperature plotted against the average across the temperature range examined, n = 3 +/- SD. The colored lipid species account for the largest abundance lipid fraction, the low abundance lipids are shown in gray. **c**. Total lipidomic remodeling employed by *M. mycoides* and Syn3B across growth temperatures; for *M. mycodes*, the results for 37-25°C and 37-30°C are shown. The overall bar height shows the total remodeling, split into remodeling magnitude of low abundance lipids (gray-colored species on Fig.6b) and of colored most abundant lipids on Fig.6b, colored part of the bar.

#### 2.6.2 Lipid class variability

To elucidate the patterns of variation that are embedded in the multitude of low abundance and low variability lipids, we return to an analysis of the lipidome in terms of aggregated lipid features, but now in the context of remodeling. By aggregating the lipidome according to class we can resolve how cholesterol varies relative to phospholipids, and how phospholipid head groups are regulated independent of acyl chain composition. *M. mycoides* and Syn3B show similar degree of variability for each class. Notably, classes that do not have a highly variable species clustering outlier (Fig.7a) exhibit comparable variability as those classes that do contain a highly variable species. This emphasizes again the key contribution to adaptation of small variations of many lipids that share a common structural feature, such as a head group. Interestingly, *M. mycoides* and Syn3B exhibit very different patterns of lipid class variability (Fig.7a). Cholesterol shows an overall decrease at lower temperatures for *M. mycoides*, whereas Syn3B cholesterol abundance increases at lower temperatures. Similarly, phospholipid head groups each vary in opposite directions when *M. mycoides* and Syn3B are compared. These patterns of variation indicate that lipid class remodeling plays a role in adaptation, and that despite having the same core genome, Syn3B exhibits a very different remodeling program from *M. mycoides*.

#### 2.6.3 Phospholipid acyl chain variability

We can further aggregate the lipidome by acyl chain features such as unsaturation (number of double bonds) and length (number of carbons), to reveal how acyl chains are remodeled independent of head group. Remodeling of lipid acyl chains is one of the primary means by which cells adapt their membrane properties to varying conditions like temperature^485411^. Increasing the degree of unsaturation (average number of double bonds per lipid) or decreasing the average chain length both lead to an increase in fluidity of the membrane that can compensate for decreased temperature.

We observed that for glycerophospholipids (GPLs: CL, PG, PC) *M. mycoides* unsaturation and chain length are relatively invariant whereas both parameters increase at lower temperature for Syn3B (Fig.7b, c (i)). It is quite remarkable that *M. mycoides* exhibited such low variation in average unsaturation over such a broad temperature range. We can dig deeper by plotting the average unsaturation and chain length of individual GPLs (Fig. 7d), which reveals that these features are varying in both *M. mycoides* and Syn3. However, when we look at the trends in average unsaturation for each phospholipid head group, we observed that PG and PC vary in opposite fashion to CL, such that the collective average change of all GPLs in *M. mycoides* is very low. This type of head group independent acyl chain remodeling has been previously reported in Gram-negative bacteria and yeast^45^, and could represent a mechanism for fine tuning membrane properties^4^.

**Figure 7.**
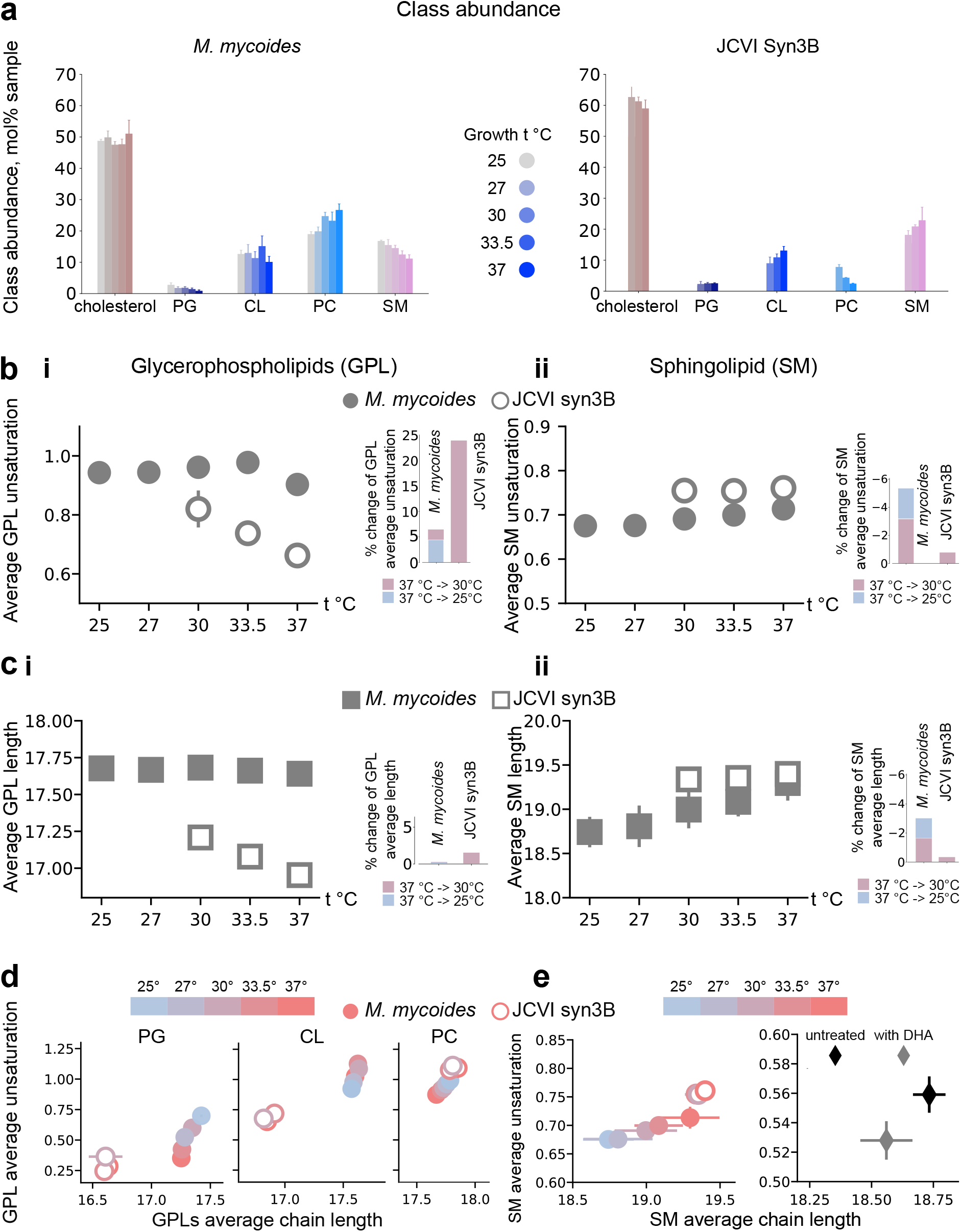
Mycoplasma features analysis reveals head-group specific acyl chain remodeling. **a.** Classes abundances at each growth temperature for *M. mycoides* and Syn3B. mean +/- SD, n = 3. **b**. Average unsaturation per acyl chain in GPLs (i) and SM (ii) against temperature: *M. mycoides* (filled circles) and Syn3B (outlined circles). mean +/- SD, n = 3. **c.** Average length per acyl chain in GPLs (i) and SM (ii) against temperature - *M. mycoides* (filled squares) and Syn3B (outlined squares). mean +/- SD, n = 3. **d**. GPLs: average chain length plotted against average unsaturation for each growth t°C. *M. mycoides* (filled circles) and Syn3B (outlined circles). mean +/- SD, n = 3. **e**. On the left: SM average chain length and unsaturations show the differences in SM features remodeling in *M. mycoides* (filled circles) and syn3B (outlined circles). mean +/- SD, n = 3. On the right: SM average features plotted against each other for RBL-derived GPMVs. n = 4 +/- SD, data adopted from Levental et.al (2016)

The patterns of sphingomyelin acyl chain remodeling were opposite of what we observed for GPLs: total average unsaturation and chain length both decreased at lower temperatures for *M. mycoides*, but were relatively invariant for Syn3B (Fig.7b,c (ii)). This suggests that for temperature adaptation, *M. mycoides* relies heavily on SM acyl chain remodeling, whereas Syn3B relies more on GPL acyl chain remodeling.

Decreasing SM chain length would increase fluidity, which would support homeoviscous adaptation at lower temperatures. The decrease in unsaturation at lower temperature, however, is contrary to the expected trend. It has been shown that the degree of unsaturation of sphingolipids can influence their interaction with cholesterol^55^, and this might be a more important factor in regulating membrane properties, than the effect of unsaturation on melting temperature. It is striking that the pattern of decreasing SM chain length and unsaturation is also observed for the plasma membrane of mammalian cells^34^ (Fig 7e). This suggests a convergent evolution in the ability to regulate SM acyl chain composition for membrane homeostasis.

### 2.7 Kinetics of lipidome adaptation

So far, we have considered the lipidome composition of cells that have been adapted to growth at a given temperature. We were also curious how the lipidome was remodeled over time following a change in temperature. To examine the kinetics of lipidome adaptation to temperature change we sample the cells for 24 hours at intervals following a decrease from 37 to 25°C (or 30°C for Syn3B), and an increase back to 37°C (Fig. 8).

**Figure 8.**
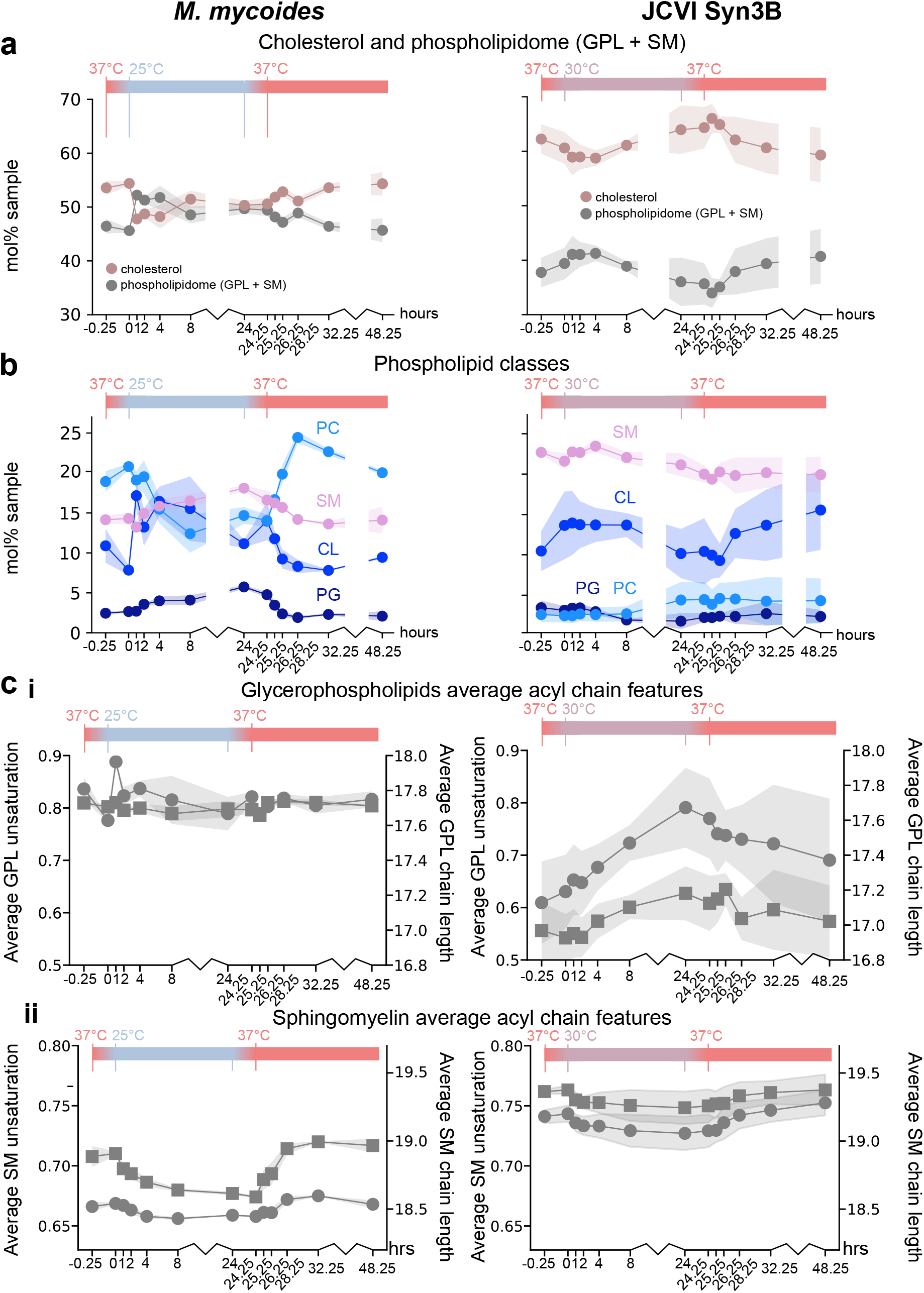
Kinetics of mycoplasma lipidome adaptation. **a.** The temperature response over time of cholesterol (rose-brown line) and other lipidome components (gray line) for *M. mycoides* (left) and Syn3B (right). The lines show the average of 3 biological replicates, bands – respective SD. The temperature dynamics with time is shown as the colored bar at the top of the plot. **b**. Phospholipid classes temperature dynamics for *M. mycoides* (left) and Syn3B (right). n = 3 +/- SD. **c**. Average unsaturation (circles) and length (squares) per acyl chain in GPLs (i) and SM (ii) dynamics with temperature in *M. mycoides* (left) and Syn3B (right). n = 3 +/- SD. The inset plot in *M. mycoides* average features shows CL average unsaturation dynamics with temperature. n = 3 +/- SD.

We again aggregate the lipidome by different features, progressing from class to acyl chain length and unsaturation. First, we considered cholesterol relative to the rest of the lipidome (Fig. 8a), which revealed unexpectedly large changes for both *M. mycoides* and Syn3B. We observed an initial large (∼7 mol%) decrease of cholesterol following a shift to lower temperature, which was relatively rapid for *M. mycoides* (∼1 hour) compared to Syn3B (∼4 hours). Subsequently, cholesterol began to return to its starting abundance. Steady-state abundance for *M. mycoides* was reached at around 8 hours, but for Syn3 cholesterol appears to still be adapting at 24 hours. Following a return to 37°C the pattern was essentially reversed, although with a smaller amplitude. *M. mycoides* appears to exhibit an overshoot at around 4 hours following the shift back to 37°C.

When plotted separately, the phospholipids (PG, CL, PC, and SM) exhibited different patterns of variation (Fig.8b). For *M. mycoides*, CL mirrored cholesterol abundance and exhibited a fast increase followed by a slower return to its starting abundance. PC decrease nearly 10 mol% over 8 hours, whilst PG and SM increased monotonically for 24 hours. The adaptation pattern was reversed following a return to 37°C with PC, CL and SM returning to staring abundances after 8 hours. Interestingly PC appears to exhibit an overshoot, at 4 hours corresponding to the cholesterol overshoot mentioned above. Syn3B showed an entirely different pattern, in which CL was the only phospholipid that changed significantly, and variations occurred gradually over both 24-hour intervals. Thus, it appears that transient cholesterol and CL homeostasis is conserved in both *M. mycoides* and Syn3B, although occurs much more rapidly in the former (Fig.8b). Additionally, Syn3B exhibits less phospholipid class adaptation than *M. mycoides*.

Next, we look at how acyl chain adaptation proceeds following temperature changes. For GPLs (CL, PG, PC) *M. mycoides* shows no significant changes in average chain length or unsaturation, except for a brief spike in unsaturation at 1 hours following a shift to 25°C (Fig.8c (i)). This spike is mostly due to a large rapid increase in CL abundance and unsaturation (Fig. 8c (i) inset). Syn3B on the other hand shows a steady monotonic rise and fall of both unsaturation and chain length during each 24-hour interval. For SM, both unsaturation and chain length adapted within 4-8 hours for both *M. mycoides* and Syn3B, with change being considerably larger for the former, consistent with our previous observations of cells adapted to lower temperatures (Fig.8c (ii)).

The adaptation patterns we observe across the lipidome were far more complex than we had anticipated and included transient changes that are lost in the analysis of cells adapted to a given growth temperature. We observed several indications of overshoot that is characteristic of biological control systems^56^. This was particularly clear for *M. mycoides* warm adaptation for PC and cholesterol. A similar overshoot behavior was recently characterized in *E. coli*, where it is proposed to play a critical role in rapidly responding to temperature fluctuations^54^. We also observe a fast and slow response to cold adaptation, that was especially pronounced in *M. mycoides*. The two-stage lipidomic response to cold shock in *M. mycoides* involves a rapid cholesterol efflux, followed by the gradual acyl chain remodeling. Such two-stage responses to temperature change have been observed in other bacteria, such as *Bacillus subtilis*^115758^, *Bacillus megaterium*^5960^ and more complex eukaryotic systems, such as carp liver^61^. For example, within minutes following a cold shock, *B. subtilis* utilizes acyl chain unsaturation as a rapid response to changes in membrane biophysical state^58^, followed by long-term changes in acyl chain branching^11^. In E.coli, the response to cold shock involves a sharp change in FabA/FaB ratio, gradually restored in long-term adaptation^54^. The presence of a two-stage adaptation in such a simple organism like M. mycoides and Syn3B might indicate that it is an essential principle of cold adaptation in living membranes as it’s performed by organisms of varying complexity with very different membrane lipid compositions.

## Discussion

In this study we have examined the ability for two simple organisms to adapt their membranes to changing temperature through lipidomic remodeling. Both *M. mycoides* and Syn3B demonstrated an ability to adjust their lipidome composition and maintain membrane fluidity across their viable temperature range. However, despite having the same core essential genome and comparable lipidome compositions, these two organisms exhibited different patterns of lipidomic remodeling. This is a remarkable demonstration of how membrane adaptation can be achieved through different patterns involving the same set of lipids, implying that the vast chemical space of the lipidome offers many divergent solutions to the same challenge. This might on the one hand cast doubt on the possibility of constraining and defining principles for lipidome adaptation. Given the complexity of lipidomes, there are an astronomical variety of ways that composition can be adjusted, and potentially much redundancy in the outcomes. On the other hand, we observe convergent patterns across species, such as SM acyl chain remodeling that is exhibited by *M. mycoides* and mammalian plasma membranes^34^ (Fig.7e). Such convergence hints at a universal, yet versatile set of principles dictating how life manages its lipids.

### Homeoviscous adaptation

Both *M. mycoides* and Syn3B exhibit homeoviscous adaptation in response to environmental temperature change, revealing that biophysical homeostasis is important even for minimal cell membranes. The concept of homeoviscous adaptation was first reported by Sinensky in the 1970s in bacterial membranes^48^ and is one of the most well-known features of living membranes^41^. Since that time, homeoviscous adaptation has been reported in membranes from microbes^62126364^ to mammals^65^, and is considered a defining characteristic of living membranes. It was reasoned, that cells maintain their membrane viscosity within an optimal range to support bioactivity^6667^. However, recent observations have brought this simple idea into question^4262^. In some bacteria, viscosity can be experimentally tuned and does not influence cell growth rates^4262^. Nonetheless, these same bacteria, naturally perform homeoviscous adaptation in response to temperature change^54^. Why would they go to all this trouble if it doesn’t affect growth? One explanation is that homeoviscous adaptation is important to allow cells to withstand large environmental changes, without crossing critical physical thresholds that would disrupt membrane function or integrity. For example, in B.subtilis, some antimicrobial peptides^68^ or a rapid cooling cause the membrane to phase separate, threatening many critical membrane functions^62^. Remarkably, there is evidence suggesting that cells tend to maintain the melting transition temperature of their membranes around 15-20 C below their growth temperature^6970^. For a soil bacterium, this could provide enough buffer to prevent crossing the membrane transition temperature during diurnal temperature cycles. Therefore, while in some organisms, homeoviscous adaptation is crucial for optimizing growth limiting processes, whereas in many organisms it may serve a subtler role in protecting the cell from crossing functionally critical physical states resulting from large environmental perturbations.

### Lipidomic remodeling

Although both organisms can remodel their lipidomes for homeoviscous adaptation to temperature change, Syn3 appears to be deficient in certain ways. At the lipid class level, Syn3B displays a remodeling pattern of phospholipid headgroups that is comparable in magnitude to *M. mycoides*, but opposite in its trend. This divergence in lipid class remodeling is coupled to an impaired ability of the minimal cell for acyl chain remodeling, especially in SM (Fig.7b,c,d,e). Interestingly, acyl chain remodeling flexibility in *M. mycoides* yields a conserved average acyl chain profile in glycerophospholipids across all temperatures. The minimal cell, Syn3B, is unable to regulate its acyl chain profile to the same extent in response to temperature fluctuations. This suggests that the simultaneous perturbation of two crucial lipidomic mechanisms – phospholipid class remodeling and fatty acid remodeling – causes Syn3B to have different average acyl chain configurations at lower temperatures. In contrast, *M. mycoides* has the distinct advantage of being able to independently regulate average acyl chain features within each phospholipid head group. This leads to the question: what are the implications of impaired lipid remodeling for Syn3B? It is conceivable that the reversed phospholipid head-group remodeling pattern exhibited by Syn3B represents a strategic effort to maximize adaptive potential within the constraints as a genetically minimal system. Alternatively, it is possible that a significant portion of the membrane sense and response mechanisms in Syn3B are compromised due to extreme genome limitations, resulting in an adaptation that is stochastic. However, their ability to at least partially maintain membrane fluidity across temperatures disfavors the latter possibility. The adaptive patterns of Syn3B accommodate both of these scenarios, underscoring the need for further investigation to unravel the fundamental lipidomic requirements for adaptation in such minimal systems.

In contrast, *M. mycoides* demonstrates a more nuanced adaptation profile, offering greater lipidomic flexibility. This leads to an intriguing question: Does the capacity of *M. mycoides* to maintain constant average acyl chain features across its lipidome, while simultaneously varying these parameters independently for each headgroup, represent a critical lipidomic mechanism for adaptation? We previously showed that varying acyl chain unsaturation has a different effect on membrane fluidity dependent on the headgroup structure^4^, and that this could be one means of fine tuning fluidity. Head group-specific remodeling may also reflect heterogenous distribution of lipids. For example, acyl chain features of lipids that are closely associated with protein transmembrane domains might require different adjustments than lipids in bulk bilayer^7172^. Furthermore, these modifications may enable the cell to concurrently fine-tune various membrane characteristics, including fluidity, permeability, and phase behavior. Additionally, the capability to independently control diverse chemically distinct components of the lipidome introduces additional degrees of freedom for membrane adaptation. Even a very restricted habitat of a simple pathogen like mycoplasma involves temporal environmental fluctuations in temperature, humidity and salinity^7374^. It is reasonable to argue that independent variation of lipidomic parameters gives the organism more room for a targeted response to different perturbations, while maintaining overall biophysical state.

### Convergent patterns of sphingomyelin remodeling across domains

Remarkably, despite their simplicity, mycoplasmas showcase patterns of lipidome remodeling that are convergent with considerably more complex organisms. *M. mycoides* coregulates SM acyl chain features, decreasing unsaturation and chain length in tandem (Fig.7e) – a pattern observed previously in mammalian plasma membrane lipidomes^34^. The precise role of sphingolipids in Mycoplasma membranes is an open question. The convergence of SM remodeling patterns in mycoplasma and mammalian membranes suggests an evolutionary conserved mechanism of utilizing sphingolipids in maintaining membrane homeostasis in environmental perturbations. Similar to cholesterol, sphingolipids are typically eukaryotic lipids, though not exclusive to Mycoplasma in the bacterial world: some prokaryotes synthesize their own sphingolipids^75^, while others utilize host sphingomyelin for adhesion and virulence^76^ . Moreover, mollicutes are not unique in their ability to scavenge sphingomyelin from the host cells – protozoan parasite *Taxoplasma gondii* is capable of its own sphingolipid synthesis, and of scavenging if from host cells^77^. Nevertheless, utilization of both cholesterol and SM by Mycoplasma showcases the unique composition of their membranes, which is in many way convergent with tremendously more complicated mammalian lipidomes. And while the mammalian membranes lipid composition is challenging to manipulate, mycoplasma membranes are tuneable^3^, and therefor offer a promising choice for further investigation of the role of sphingomyelin in membrane adaptation.

### Mechanisms of lipid uptake and remodeling in mycoplasmas

Despite the long-standing research interest in mycoplasmas^78^ as a potent minimal membrane model system, our understanding of how these parasites take up and remodel their lipids remains elusive. The incorporation of free and esterified cholesterol is a staple feature of parasitic mycoplasmas - the saprophytic strains, such as *A. laidlawii*, do not require sterol for growth^1779^. Free or esterified cholesterol cannot be chemically remodeled by mycoplasmas^79^ – sterols remain an exogenous membrane component, however, mycoplasmas can control sterol content of their membranes^80^, which is also confirmed by our experiments (fig.8). Early studies on the role of sterols in mycoplasma membranes speculated that cholesterol serves a primary role as a membrane rigidifying component and compensates for lack of a cell wall and limited phospholipid synthesis^81^. Indeed, one of the explanations for the two-stage adaptation to temperature change (Fig.8) might be directly derived from the role of sterols in mycoplasma membrane structural integrity^82^ and biophysical properties. Cholesterol is a mediator of membrane fluidity in mycoplasma^83^, and was shown to reduce osmotic sensitivity of mycoplasma cells^84^. It might therefore be possible that mycoplasmas employ cholesterol for rapid adaptation to environmental stressors but do not allow long-term fluctuations in sterol abundance. The precise mechanism of cholesterol uptake and regulation in mycoplasmas is, however, still unknown. It’s been shown that isolated mycoplasma membranes retain the capacity for cholesterol uptake^80^, and the hypothesis is that it is partially a process of physical adsorption^80^. Furthermore, it is known that sterol uptake is correlated with phospholipid abundance in membranes, as cultures with experimentally modified phospholipid concentrations experience a consecutive change in sterol amounts^85^. This observation allows to hypothesize that phospholipid binding is a crucial step in cholesterol homeostasis in mycoplasma membranes. Our experiments of time-temperature adaptation suggest a presence of cholesterol efflux mechanism that allows for a rapid sterol drop in response to temperature decline. While such an efflux mechanism is directly coupled to phospholipid abundance in the membrane, the physicochemical interactions in the absence of active transport are unlikely to explain a rapid change in cholesterol abundance. It is therefore plausible that an active sense-and-response mechanism is present in mycoplasma cells that detects changes in membrane biophysical properties and initiates short-term cholesterol depletion. The gradual restoration of cholesterol levels in the course of long-term adaptation may involve both active (sense-and-response) and passive (physical adsorption) mechanisms.

In contrast to cholesterol uptake, early experiments with isolated mycoplasma membranes showed that no significant phospholipid uptake occurs in the absence of cellular metabolism^8086^, which suggested active mechanisms of phospholipid uptake. The experimental evidence suggests that mycoplasmas retain enzymatic machinery to modify acyl chain composition of phosphatidylcholine^86^, but not sphingomyelin^8687^ – two major external phospholipid classes, taken from the serum^86^. Very little is known about precise mechanisms of phospholipid uptake and homeostasis in mycoplasmas. A very recent study on pathogenic *Mycoplasma pneumoniae* sheds some light on the mechanism that mycoplasma employs to scavenge lipids from the host^88^: a newly-discovered essential surface protein P116 directly interacts with HDLs in the serum and takes up cholesterol, PC and SM. Further, it is known that parasitic strains are incapable of acyl chain synthesis^89^, relying on the external inputs to perform internal lipid synthesis and modification. However, the existing experimental evidence gives no information about acyl chain recognition and how the available acyl chain pool is distributed among different phospholipid classes. The previous experimental evidence and our results suggest that there is likely a combination of active mechanisms that facilitate cholesterol and phospholipid uptake and remodeling. However, the possibility of partial non-selective phospholipid uptake cannot be excluded.

### Outlook

Looking forward, a critical step in identifying the mechanisms of lipid remodeling in mycoplasma will be to challenge the organism with a defined restriction in lipid input. Animal serum, currently used as a lipid supplement, provides a complex and varied array of dietary lipids, enabling mycoplasma to synthesize an optimally adaptable membrane from a structurally diverse pool of lipids. As a result, the role of internal lipid remodeling in these optimal dietary conditions is challenging to decipher. By taking control over the lipid diet of mycoplasmas, it will be possible to systematically examine which lipidomic remodeling patterns are essential for mycoplasma cellular viability.

In summary, we have juxtaposed a simplistic parasitic bacterium, *M. mycoides*, with its synthetic counterpart, JCVI Syn3B, to unravel the principles governing membrane lipidomic adaptation in response to environmental temperature shifts. Our results show that the Minimal Cell retains an ability for active lipidomic remodeling and homeoviscous adaptation, despite an impaired capacity for head-group specific acyl chain regulation observed in *M. mycoides*. Our findings provide the first evidence for lipidomic remodeling and membrane adaptation in a minimal living organism and reveal the value of mycopalsma as a minimal living membrane model system. Lipidomic and environmental perturbations of *M. mycoides* and Syn3B offer a great potential for deciphering the minimal requirements for a responsive bioactive cell membrane.

## Acknowledgements

The authors wish to thank Sáenz Lab members Tomasz Czerniak, Isaac G. Justice, Ha Ngoc Anh Nguyen, as well as Ilya Levental and Kandice Levental for discussions and feedback. In addition, the authors wish to thank the colleagues from J. Craig Venter Institute (JCVI) – John Glass and Kim Wise – for discussions and feedback, including manuscript comments. We thank Telesis Bio, Inc. for providing us with the bacterial strain JCVI-syn3B. We also thank Lipotype GmbH for generous assistance with lipidomic samples preparation and analysis. Finally, we are grateful to Andrey Klymchenko and Dmytro Danylchuk from the Faculté de Pharmacie, Université de Strasbourg for providing us with pro12A for lipid order measurements.This work was supported by the B CUBE of the TU Dresden, a German Federal Ministry of Education and Research BMBF grant (to J.S., project 03Z22EN12), and a VW Foundation ‘‘Life ’’grant (to J.S., project 93090).

## Bacterial strains

### M. mycoides subsp. capri strain GM 12

JCVI-Syn3B

Both bactetrial strains were received from J. Craig Venter Institute (JCVI) (La Jolla, California, USA)

### Chemicals

Both mycoplasma strains were cultivated in SP4 growth medium, supplemented with CMRL and 17% FBS. The original SP4 recipe was first developed by Tully et.al in 1970s ^62^; later, colleagues from J.Craig Venter Institute supplemented SP4 with extra nutrients to support viability of the minimal cell^19^. The detailed SP4 composition is listed in Table 1 in Supplementary Materials.

Pro12A dye was a courtesy of Klymchenko Lab^49^.

### Mycoplasma Growth in Parallel DasGIP Bioreactor System

*Bioreactor Setup*: A 4x parallel DasGIP bioreactor system was employed for the cultivation of *M. mycoides* and Syn3B for all lipidomics experiments. Each bioreactor vessel was sterilized by autoclaving with 500 mL of SP4 media supplemented with fetal bovine serum as a lipid source. Media input reservoirs and waste reservoirs were connected to the reactors using a sterile tube welder to ensure aseptic conditions.

*Turbidostatic Control*: The bioreactor system was operated in turbidostatic mode, aiming to maintain a constant cell density and thereby maximizing the growth rate of *M. mycoides* and Syn3B. Optical density (OD) measurements were taken at a wavelength of 600 nm, serving as the primary feedback control. 25 ml of batch culture in mid-exponential growth phase was pelleted at 12000g for 2 min and resuspended in 3 ml of fresh growth medium (SP4 + FBS). The cell suspension was used to inoculate bioreactors.

*Instrument Calibration*: Probes, including those for OD measurements, were calibrated using a procedure integrated into the DasGIP software.

*System Regulations*: The pH, O2 concentration, and temperature within each bioreactor vessel were automatically monitored and regulated by the DasGIP reactor system software. Specific set points for these parameters were set at pH 7, O2 at 30% of atmospheric, and temperatures ranging from 25 to 37°C. *Gas Flow and Agitation*: Reactors were continuously gassed with air at a flow rate of 3-20 sL/h. Agitation was achieved using an impeller, with speeds varying between 300 to 500 rpm, depending on the real-time requirements of the cell culture.

*Media Composition and Anti-foam Agent*: The growth media was supplemented with 4 ml of anti-foam agent (Roth Art.-Nr. 0865.1, 1:5000 dilution) to prevent excessive foaming during the cultivation process. *Level Control*: A level sensor was used to ensure the reactor volume remained constant. Any deviations from the set volume triggered adjustments in media input/output rates by the system.

*Sampling and Monitoring*: Mycoplasma cells were sampled manually at various intervals throughout the cultivation period by withdrawing media using an external peristaltic pump. These samples were taken to validate the turbidostatic control method and to ensure that cells remained in their exponential growth phase. Cells were streaked out on SP4 and LB solid agar plates, and inspected by phase contrast microscopy to ensure no contaminating organisms were present.

### Mycoplasma membrane separation

*M. mycoides* and Syn3B membranes were separated using sucrose gradient. Cell cultures were grown in batch in 500 ml glass flasks and harvested at early-exponential phase (OD 600 0.2-0.3 for *M. mycoides*, OD 600 0.07-0.15 for Syn3B) and washed twice for 10 min at 7000g in mycoplasma wash buffer (200 mM NaCl, 1% glucose, 25 mM HEPES, pH 7.0). Next, cell suspension was lyzed using Avestin Emulsiflex- B15 and the lysate was placed on ice. Lyzed samples were spinned at 10000g for 7 min at 4°C to remove remaining cellular debris. Lysates were loaded onto 65-3010 % (w/v) sucrose gradient and ultracentrifuged overnight (at least 8 hours) at 4°C, 250000 g. Membrane fractions were collected in separate tubes and washed twice to remove remaining sucrose (4°C, 70000g 1 hour each time). Finally membrane fractions were resuspended in 450 uL mycoplasma wash buffer and kept at -20°C analysis. Phosphate assay was used to determine precise lipid concentration of membrane fractions. 50 umols of membrane fractions in mycoplasma wash buffer were submitted for shotgun lipidomics.

### Determination of membrane lipidome composition using shotgun lipidomics

#### 1. Whole cell sample preparation for lipidomic analysis

Bacterial samples were harvested at designated time points from the continuous cell culture in the bioreactor system and washed twice in mycoplasma wash buffer (200 mM NaCl, 25 mM HEPES, 1% glucose, pH 7.0) to remove traces of the growth medium at their growth temperature (21000g, 2 min centrifuging steps). Washed cell pellet was resuspended in 150 uL PBS and flash-frozen in liquid nitrogen.

#### 2. SP4 + FBS sample preparation for lipidomic analysis

88.2 uL of SP4 + FBS were diluted in 661.8 uL milliQ H2O and submitted for lipidomics analysis in 4 replicates

#### 3. Lipid extraction for mass spectrometry lipidomics

Lipids were extracted using a two-step chloroform/methanol procedure ^6^. Samples were spiked with internal lipid standard mixture containing: cardiolipin 14:0/14:0/14:0/14:0 (CL), ceramide 18:1;2/17:0 (Cer), diacylglycerol 17:0/17:0 (DAG), hexosylceramide 18:1;2/12:0 (HexCer), lyso-phosphatidate 17:0 (LPA), lyso-phosphatidylcholine 12:0 (LPC), lyso-phosphatidylethanolamine 17:1 (LPE), lyso- phosphatidylglycerol 17:1 (LPG), lyso-phosphatidylinositol 17:1 (LPI), lyso-phosphatidylserine 17:1 (LPS), phosphatidate 17:0/17:0 (PA), phosphatidylcholine 17:0/17:0 (PC), phosphatidylethanolamine 17:0/17:0 (PE), phosphatidylglycerol 17:0/17:0 (PG), phosphatidylinositol 16:0/16:0 (PI), phosphatidylserine 17:0/17:0 (PS), cholesterol ester 20:0 (CE), sphingomyelin 18:1;2/12:0;0 (SM), triacylglycerol 17:0/17:0/17:0 (TAG) and cholesterol D6 (Chol). After extraction, the organic phase was transferred to an infusion plate and dried in a speed vacuum concentrator. 1st step dry extract was re-suspended in 7.5 mM ammonium acetate in chloroform/methanol/propanol (1:2:4, V:V:V) and 2nd step dry extract in 33% ethanol solution of methylamine in chloroform/methanol (0.003:5:1; V:V:V). All liquid handling steps were performed using Hamilton Robotics STARlet robotic platform with the Anti Droplet Control feature for organic solvents pipetting.

#### 4. MS data acquisition

Samples were analyzed by direct infusion on a QExactive mass spectrometer (Thermo Scientific) equipped with a TriVersa NanoMate ion source (Advion Biosciences). Samples were analyzed in both positive and negative ion modes with a resolution of Rm/z=200=280000 for MS (species structural resolution; see Table 2 in Supplementary Materials) and Rm/z=200=17500 for MSMS (subspecies structural resolution; see Table 2) experiments, in a single acquisition. MSMS was triggered by an inclusion list encompassing corresponding MS mass ranges scanned in 1 Da increments ^9^. Both MS and MSMS data were combined to monitor CE, DAG and TAG ions as ammonium adducts; PC, PC O-, as acetate adducts; and CL, PA, PE, PE O-, PG, PI and PS as deprotonated anions. MS only was used to monitor LPA, LPE, LPE O-, LPI and LPS as deprotonated anions; Cer, HexCer, SM, LPC and LPC O- as acetate adducts and cholesterol as ammonium adduct of an acetylated derivative ^91^.

### Lipid extraction and MS data acquisition have been performed by Lipotype GmbH

#### 5. Lipidomic data filtering

Each sample’s data was obtained in 3 biological replicates and each lipid species is reported as mol% of the total sample for each temperature and each bacterial strain (i.e, all lipid species sum up to 100% for each replicate). Prior data analysis, storage lipids (cholesteryl esters and TAGs) were excluded and the remaining lipid abundances recalculated to 100 mol%. To filter the data, the average of 3 replicates was calculated for each lipid species in the sample. The average was sorted from the most to the least abundant species and the respective cumulative sum was calculated. 2% of the least abundant species in cumulative sum were filtered out. The remaining 98% of the most abundant species were considered for further analysis. The filtered data were further resummarized to 100%. Other published lipidomes analyzed in this study were also filtered using the 98% filtering threshold.

### Published lipidomes analyzed in this study

1. *M. extorquens* filtered lipidome was taken from ^4^; the following conditions were considered for further analysis (in biological triplicates):

a. Growth stages: early-, mid-, late-exponential and stationary (harvested at 30°C)
b. Growth temperatures: 4°C, 13°C, 20°C (harvested at early-exponential growth stage)
2. *S. cerevisiae* whole cell lipidomes were taken from^5^. The following conditions were considered:

a. Growth stages: early (OD 1)-, mid (OD 3.5)-, late-exponential (OD 6) and stationary phase
b. Growth temperatures: 15°C, 24°C, 30°C, 37°C
3. *S. cerevisiae* vacuole lipidomes from^7^. Conditions considered:

a. Exponential growth stage (4 replicates)
b. Stationary growth stage (4 replicates)
4. Plasma membrane GPMVs lipidomic data was taken from ^34^:

a. Rat basophilic leukemia cells sample #1, untreated and treated with DHA
b. Rat basophilic leukemia cells sample #2, untreated and treated with DHA

### Lipidomic data analysis

1. **Data grouping**. For various purposes of lipidomic analysis, the filtered list of lipidomic species was grouped by different structural properties of lipids: structural category, head group, length or unsaturation, in most cases yielding the smaller subset of lipidomic entries. Therefore the resulting subset was resummarized to 100 mol% prior subsequent analysis.
2. **Principal component analysis.** Filtered mycoplasma lipidomic datasets in triplicates were considered for PCA. Python sklearn package was used for analysis. For Fig.2a PCA, phosphatidylglycerol (PG) and cardiolipin (CL) species were first excluded from datasets and the remaining species were resummarized to 100 mol%.
3. **Lipidomic remodeling.** Unless stated otherwise, lipidomic remodeling of a given lipidomic subset is a sum of absolute changes of species/features averages across 2 growth temperatures, as indicated by the following formula:

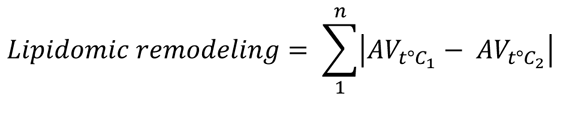

Where:

AV = species average abundance at a given temperature,

n = total number of entries in a given lipidomic subset.

**Total lipidomic remodeling** refers to the remodeling magnitude of all species in a lipidomic sample across two different growth temperatures. Standard deviations for lipidomic remodeling were propagated using pooled standard deviation.

4. Fig.6

a. Maximum curvature of lipid abundance curves was determined by kneed python package (method published by Satopaa et.al^53^)
5. **Fig**.7:

a. for class variability, standard deviation for class abundances across temperatures is considered as a readout.
b. **Average features**. Average chain length and unsaturation for each phospholipid were determined from species features according to the acyl chain number in the following way. Using the following examples:

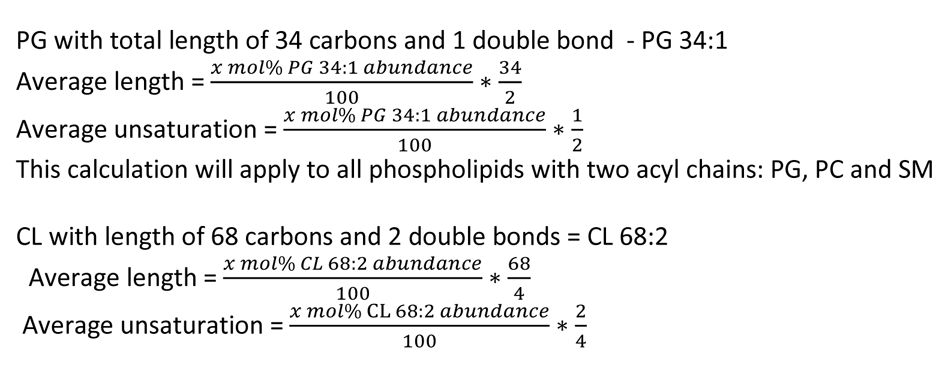

6. **Percentage change of average features**. To calculate percentage change of average features at lower temperature relative to 37°C growth temperature, the following method was used:

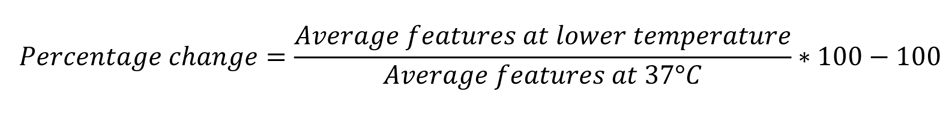

### Bacterial growth quantification using cell culture absorbance

To quantify bacterial growth at different growth temperatures, batch bacterial cultures were setup in glass 100 ml flasks to manually monitor bacterial absorbances at 600 nm over the entire growth range at each temperature. The absorbances were recorded using DeNovix DS-11 FX+ spectrophotometer/fluorometer and were corrected for the respective blank growth medium. To calculate bacterial growth rates, the slope of the linear part of the log-linear bacterial absorbance-time curves was calculated. Next, bacterial absorbances were corrected for the starting measurement at t = 0 and the maximal optical density value was taken as the maximum cellular density.

### Membrane lipid order determination using Pro12A probe

Batch bacterial cultures in glass 100 ml flasks were started in biological triplicates at each of the growth temperatures. Cells were harvested at early exponential growth state, (OD 600 nm 0.2-0.29 for *M. mycoides* and 0.05-0.15 for JCVI syn3B), washed twice (4000g, 5 min) and resuspended to final OD 0.2 in mycoplasma wash buffer. Pro12A dye was diluted to 160 uM in DMSO. 0.5 uL/200 uL cell culture was added to each well of the 96-well plate to yield final 0.4 uM dye concentration. Washed bacterial cells in mycoplasma buffer were added on top of the dye to each well. The measurement plate with labelled bacterial samples was incubated at a growth temperature for 20 min, fluorescence recorded at 440 nm and 490 nm, with 20 nm bandwidth, following excitation at 360 nm +/- 10 nm.

Pro12A general polarization was calculated according to the formula:

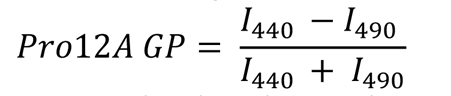

Measurements were carried out in Tecan Spark multimode microplate reader.

To ensure measurement reproducibility, restraining the harvesting window of cellular growth state was crucial. All consumables, wash buffer, Tecan Spark and the centrifuge were preheated to the bacterial growth temperature for at least 1 hour prior the measurement. *M. mycoides* cells were passaged twice a day when kept in 30-37°C growth range, daily for 27°C and once every 2 days for 25°C cells to exclude cell ageing effects on the fluorescent measurement. JCVI Syn3B cells were passaged daily. After washing steps, cellular density was estimated in the buffer to yield the final density at OD 0.2 to ensure constant dye/cell density ratio.

To determine statistical significance in Pro12A readout difference across temperatures, Mann-Whitney ranking test was used on GP values.

### Liposome preparation for Pro12A assay

POPC and DPPC lipids were purchased from Avanti as chloroform solutions. Cholesterol was obtained from Sigma Aldrich in powder form. Phospholipid concentration was determined with phosphate assay. The following lipid mixtures were prepared, with total concentration of 5 mM:

1. Cholesterol:POPC 1:1
2. Cholesterol:DPPC 1:1

Lipids were mixed in solvent-proof 2 ml Eppendorf tube and chloroform was evaporated in vacuum concentrator (Christ RVC 2-25 CDplus). Dry lipid films were rehydrated in mycoplasma wash buffer (200 mM NaCl, 25 mM HEPES, 1% glucose, pH 7.0) and incubated at 55°C for 30 min, 1000 rpm shaking. Lipid mixtures were then sonicated in the sonic bath for 5 min. Finally, 10 freeze-thaw cycles were performed on all mixtures and liposomes were stored at -20°C. Liposomes were prepared in triplicates.

For Pro12A staining the same protocol was used as for mycoplasma cells. Liposomes were diluted in mycoplasma wash buffer to 100 uM concentration and stained with 0.25 mol% dye.

