## Supplementary Figures for "From hot to cold: dissecting lipidome adaptation in *Mycoplasma mycoides* and the Minimal Cell JCVI-Syn3B"

**a**

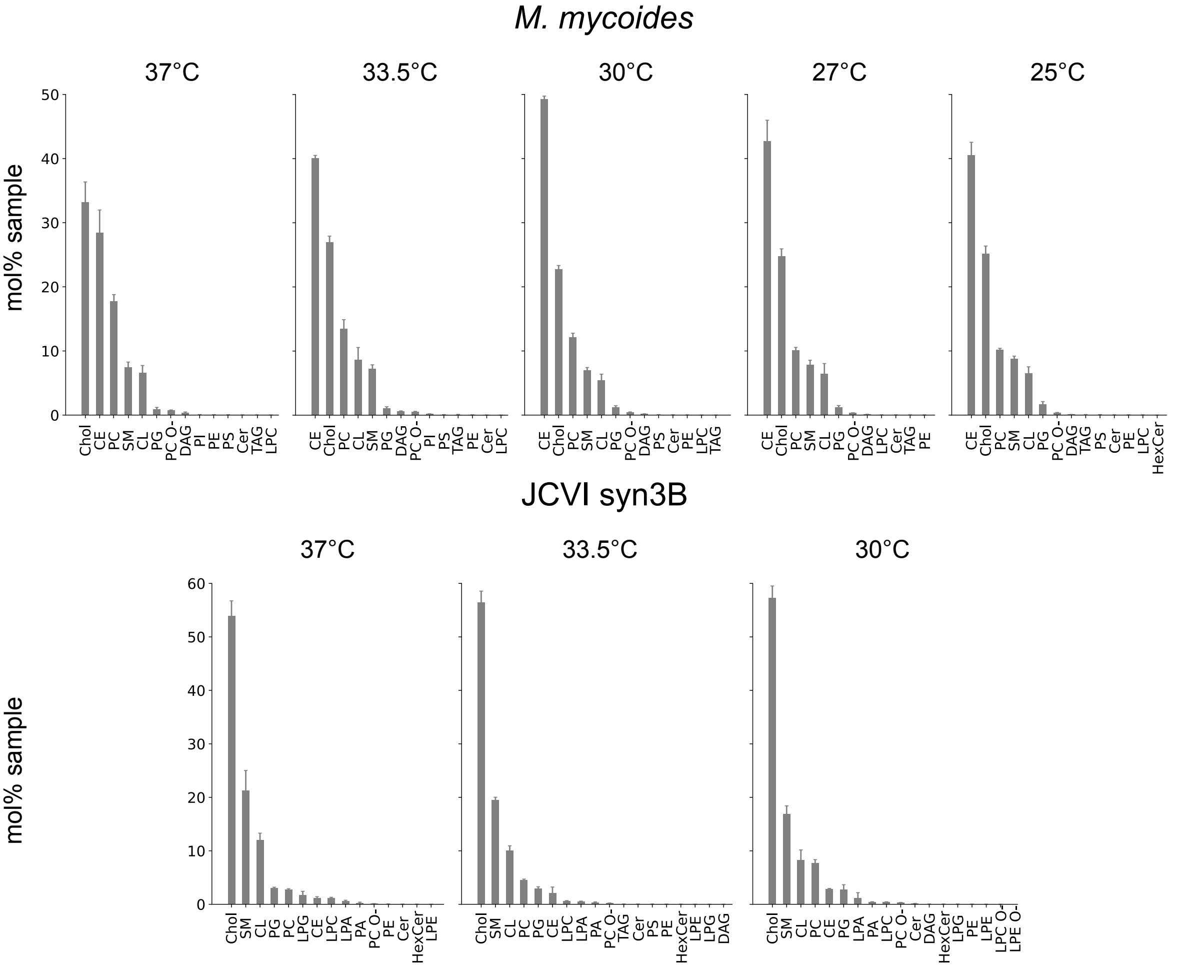


**b**

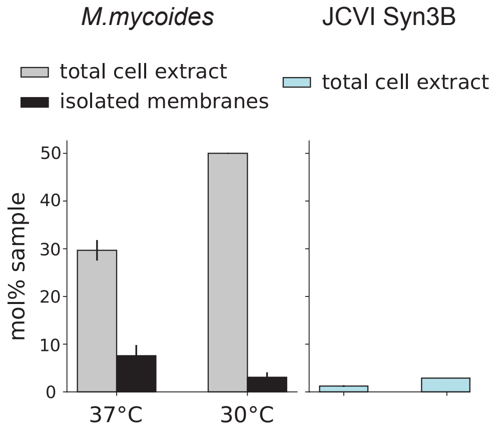


**Supplementary Figure 1.a.** All lipid classes present in unfiltered mycoplasma lipidomes at all growth temperatures. *M. mycoides* (top) and JCVI Syn3B (bottom).n= 3 +/- SD **b**. Mol% abundance of cholesteryl esters in mycoplasma total cell lipid extracts and isolated membranes, collected from cells, grown at 37°C and30°C (mean +/- SD, n = 3).

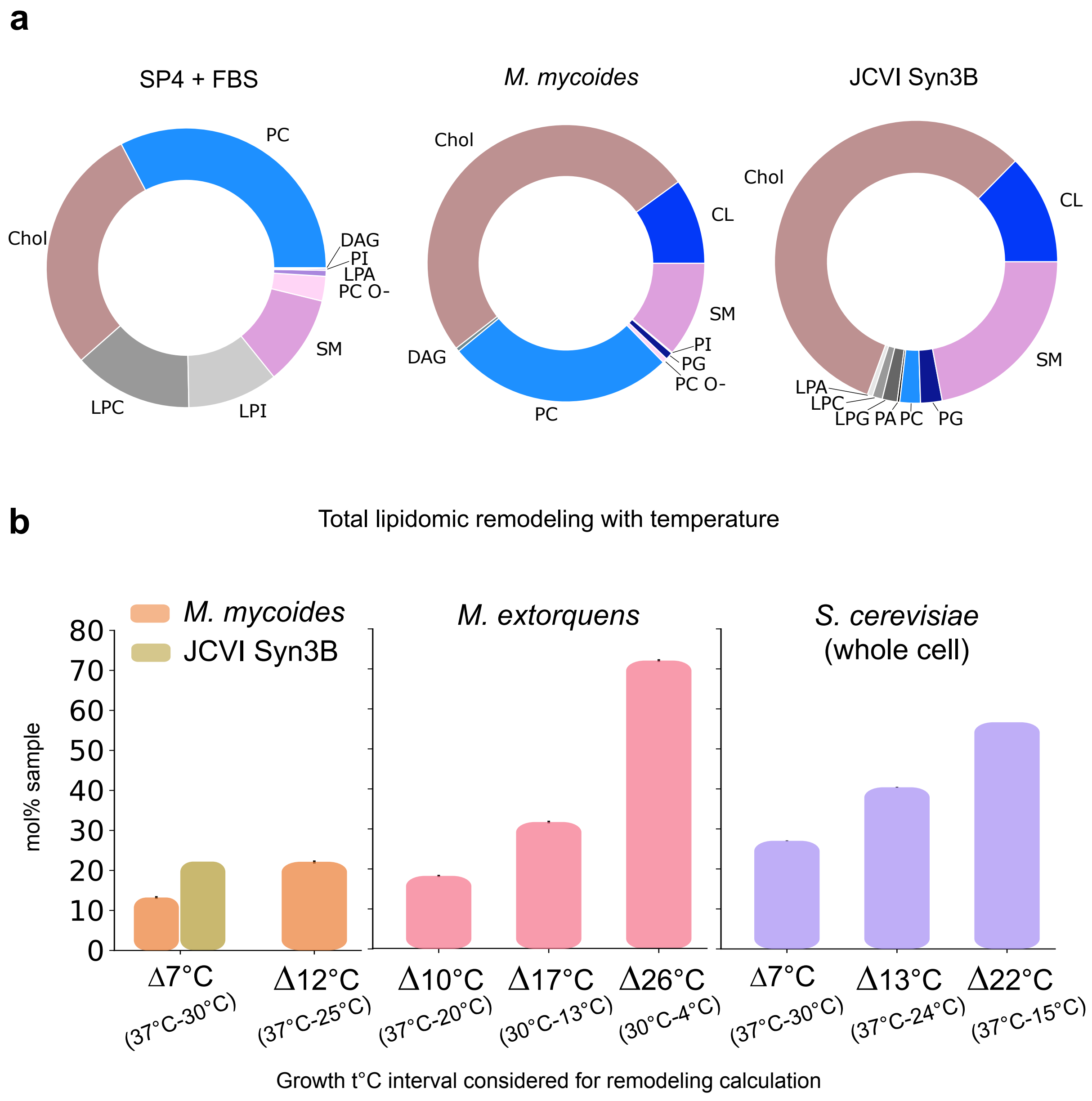


**Supplementary Figure 2. a.** Lipid classes in filtered lipidomes of growth medium (with FBS) and mycoplasma at 37°C. All storage lipids excluded. Pie chart shows average abundance, n = 3. **b.** Total lipidomic remodeling with temperature, shown for those organisms, where temperature conditions were available. Scatter markers show the average remodelig across all temperatures, with respective standard deviations.
